# Assembly of skin microbiomes is more neutral than gut microbiomes in multiple animal species

**DOI:** 10.1101/2025.07.16.665179

**Authors:** Killian D. Campbell, Brendan J.M. Bohannan, Karen L. Adair

## Abstract

The gut and external tissues of most animals are colonized by communities of microorganisms that can influence the health, development, and fitness of the host. The composition of these communities can vary greatly between individuals within a host species, and both selective factors (e.g. host immune response) and neutral processes (e.g. random loss of microbial cells) have been shown to contribute to this variation. While it is known that microbiome composition differs between tissues within an individual host, less is known about the ecological processes that underline these differences. To address this, we investigated whether the contribution of neutral ecological processes to microbiome assembly differs between external (skin and scale) and internal (gut) host tissues for a diverse panel of animal hosts. To do this, we fit a neutral ecological model to microbial communities from external and internal tissues across a variety of animal hosts. Strikingly, we discovered that the neutral model was equally or better fit to skin or scale microbial communities across all hosts, suggesting that neutral processes play a larger role in the assembly of skin or scale microbiomes compared to gut microbiomes. Furthermore, we observed that this trend is robust to different definitions of the metacommunity (i.e. the microbial taxa available to colonize a host). Finally, we leveraged a simulation framework to compare the model fits of empirical vs simulated microbial communities. We found that neutral model fits to empirical communities can differ from simulated communities, emphasizing the importance of temporal sampling in profiling animal microbiomes.

## Importance

Animal microbiomes are complex assemblages of microbial cells that influence a wide variety of host phenotypes. Despite their importance we lack a thorough understanding of the processes that guide the formation of microbiomes (i.e. microbiome assembly). Understanding how microbiomes assemble is essential to managing microbiomes for host health, conservation, and other goals. Our work highlights the relatively underappreciated role of neutral ecological processes (the random loss or gain of microbial cells) in the assembly of animal microbiomes. We document a potentially general trend: the microbiomes of external tissues (i.e. skin or scales) tend to be more neutrally assembled than those of internal tissues (i.e. guts). This observation suggests that the commonly reported differences in microbiome composition of external and internal animal tissues may be due in part to different assembly processes. Our work also highlights the dynamic nature of microbiomes, and the importance of longitudinal sampling when studying animal microbiomes.

## Observation

Most animals are colonized by communities of microorganisms that influence their development, health, and fitness (1,2). In healthy individuals, these host-associated microbial communities, or microbiomes, are predominantly found in the gastrointestinal tract, respiratory system, and external tissues (e.g. skin or scales; refs). It is clear that microbiome composition is strongly influenced by body site (3-6) but the ecological processes that underlie this variation remain unclear.

Two main types of processes contribute to the assembly of a microbiome over relatively short time scales. Selective processes are the result of fitness differences among members of a community. Examples of selective factors include variation in oxygen tolerance, ability to grow on specific carbon sources, and competitive or mutualistic interactions. In contrast, ecological drift and passive dispersal are considered ecologically “neutral” (i.e. they do not vary consistently across species) and are often modelled as random processes (7,8). Understanding the relative roles of selective and neutral processes in microbiome assembly is essential to managing microbiomes for host health, conservation, and other goals (8).

Here we investigate whether the contributions of selective and neutral processes differ between external (e.g. skin or scales) and internal (i.e. gastrointestinal tract) host tissues, and whether these patterns are consistent across host taxa. We identified 16 published datasets that used 16S rRNA gene amplicon sequencing to characterize both the external and gut microbiomes of at least 20 individuals in a set of diverse animal hosts, including humans (Supplemental table 1). For each study, the raw sequencing reads were acquired as fastq files from the NCBI Sequence Read Archive or other publicly available database and processed with the Qiime2 platform v2020.2 (9). Amplicon sequence variants (ASVs) were resolved with the dada2 pipeline (10) and classified with a naïve Bayes classifier pre-trained on the Silva 138 SSU database (11-13). Reads for all individuals were rarefied to a uniform read depth matching the lowest read depth from a given individual in the study using the vegan package in R (14). To determine the relative contribution of neutral processes to microbiome assembly, we assessed fit of Sloan Neutral Community Model for Prokaryotes (15) to the distribution of microbial taxa in these datasets as previously described (8).

Applying a metacommunity theoretical framework, the SNCM predicts the relationship between the proportion of host individuals (local communities) in which a microbial taxon occurs and its mean relative abundance across all hosts (the metacommunity). In this model, which assumes equal rates of per-capita growth and death, taxa that are only present in a few hosts are likely to be lost due to ecological drift while abundant taxa are most likely to spread among hosts due to passive dispersal. The model is fit to the observed data with the single free parameter *m*, the probability that a random loss will be replaced by reproduction from the same microbial community versus dispersal from another host (8,16). Microbial taxa that fell within a 95% confidence interval around the model prediction were considered to follow neutral assembly processes as described previously (8).

We first compared the fit of the neutral model for microbial taxa (ASVs) in gut and skin or scale tissues separately for each host species with the metacommunity restricted to samples of the same tissue type. Model fit was assessed by the root mean squared error (rmse) metric with lesser rmse values indicating a better fit of the SNCM to the data. Across all animal hosts, skin or scale microbiomes were equally or better fit to the neutral model compared to gut microbiomes (Fig 1A). These results suggest that overall, the assembly of microbiomes associated with skin or scale tissues tend to be more governed by neutral processes than gut microbiomes. However, the strength of this pattern varied among host taxonomic groups; for example, the fit of the neutral model did not differ between skin or scale and gut tissues for the fish datasets considered in this study, while skin microbiomes tended to be a better fit to the neutral model than gut microbiomes for mammal hosts.

**Figure 1:**
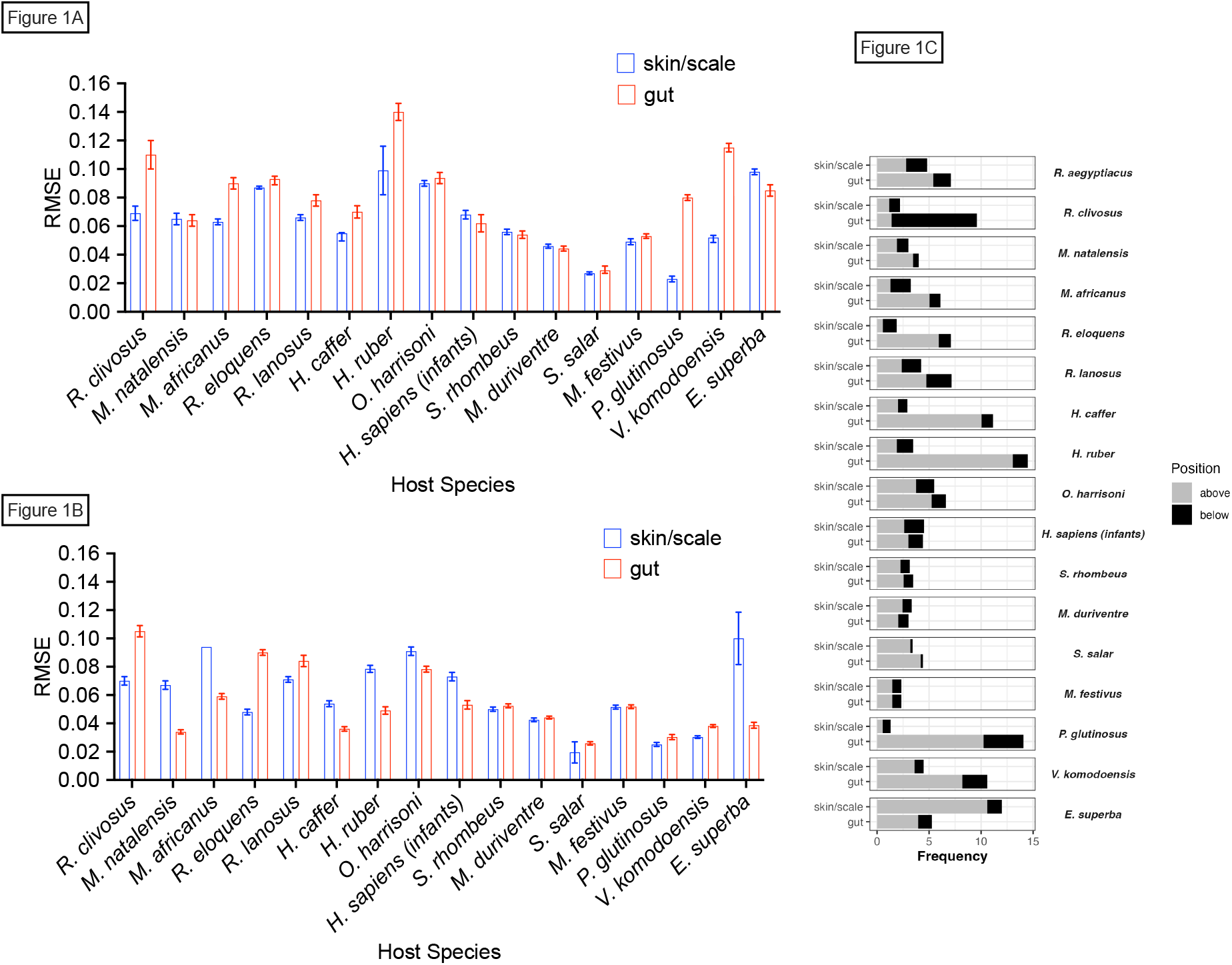
A) Neutral model fits of empirical microbial communities for both skin/scale and gut microbial communities across animal hosts. B) Neutral model fits of simulated microbial communities for both skin and scale and gut microbial communities based on community metrics obtained from corresponding empirical data. Error bars for both A and B represent 95% confidence intervals based on bootstrapping of data. C) Proportion of microbial taxa that fell outside of neutral model predictions for skin/scale and gut communities.

Next, we sought to compare the neutral model fits of empirical communities with simulated microbial communities (Fig 1B). Simulating microbial communities allowed us to obtain a steady-state level of neutrality against which we could compare our empirical communities. This is crucial, as neutrality levels may vary over time. For example, variations in neutrality are sensitive to ‘selective’ factors that may be present at varying intensities at different points in time, including changes in host immunological state due to immune system development or pathogen exposure (17,18). We leveraged an existing simulation (16) framework to generate completely neutral communities that had the same number of hosts, microbial diversity and migration parameter ‘m’ as the model fits from each empirical dataset. Simulating communities recapitulated the trends in neutrality observed in the empirical communities. However, the model was not uniformly better fit to simulated communities, indicating that microbial community composition can vary over time, and that longitudinal sampling may be necessary to fully understand microbiome assembly (Figs 1A and 1B).

To better understand the factors that contribute to the observed patterns of neutrality in skin/scale vs. gut communities, we looked for patterns in the percentage of microbial taxa that fell outside of neutral model predictions across the different hosts for each tissue type. Taxa that occur more or less frequently than neutral model predictions are thought to be non-neutrally distributed and therefore may be subject to selective processes (Fig 1C). For example, taxa that occur in more individual hosts than predicted by the neutral model (above the neutral model prediction) may be adapted to persist in the host environment or selected for by the host. In contrast, taxa below the neutral model prediction (i.e. detected in fewer samples than predicted by the SNCM) could be particularly dispersal limited or may be selected against by the host immune system. For nearly all the datasets considered in this study, the majority of the ‘non-neutral’ ASVs fell into the “above partition”, suggesting that these taxa are particularly host-adapted (Figure 1C). For the few host species where the SNCM was a better fit for the gut microbiome than the skin or scale microbiome, this was also primarily due to a higher proportion of taxa in the above partition, suggesting that for a minority of host species, the gut environment is favourable to more microbial taxa than the skin or scale environment.

Finally, we assessed whether changing the definition of the metacommunity to include microbes from both the skin or scale tissues and gut tissues of a given species influenced the fit of the neutral model. Microbes have been shown to disperse among body sites within a host individual, which suggests that a less tissue-specific definition of the metacommunity may be more appropriate (19-21). Overall, the observation that assembly of gut microbiomes tended to be equally or less neutral than the assembly of microbiomes in skin or scale tissues is consistent whether the metacommunity is restricted to a particular tissue type or includes both tissue types (Figure 2). However, there were some exceptions (e.g. *E. superba*), suggesting that careful consideration of the source and sink communities is essential for an accurate assessment of the contribution of neutral process to the assembly of host-associated microbiomes.

**Figure 2:**
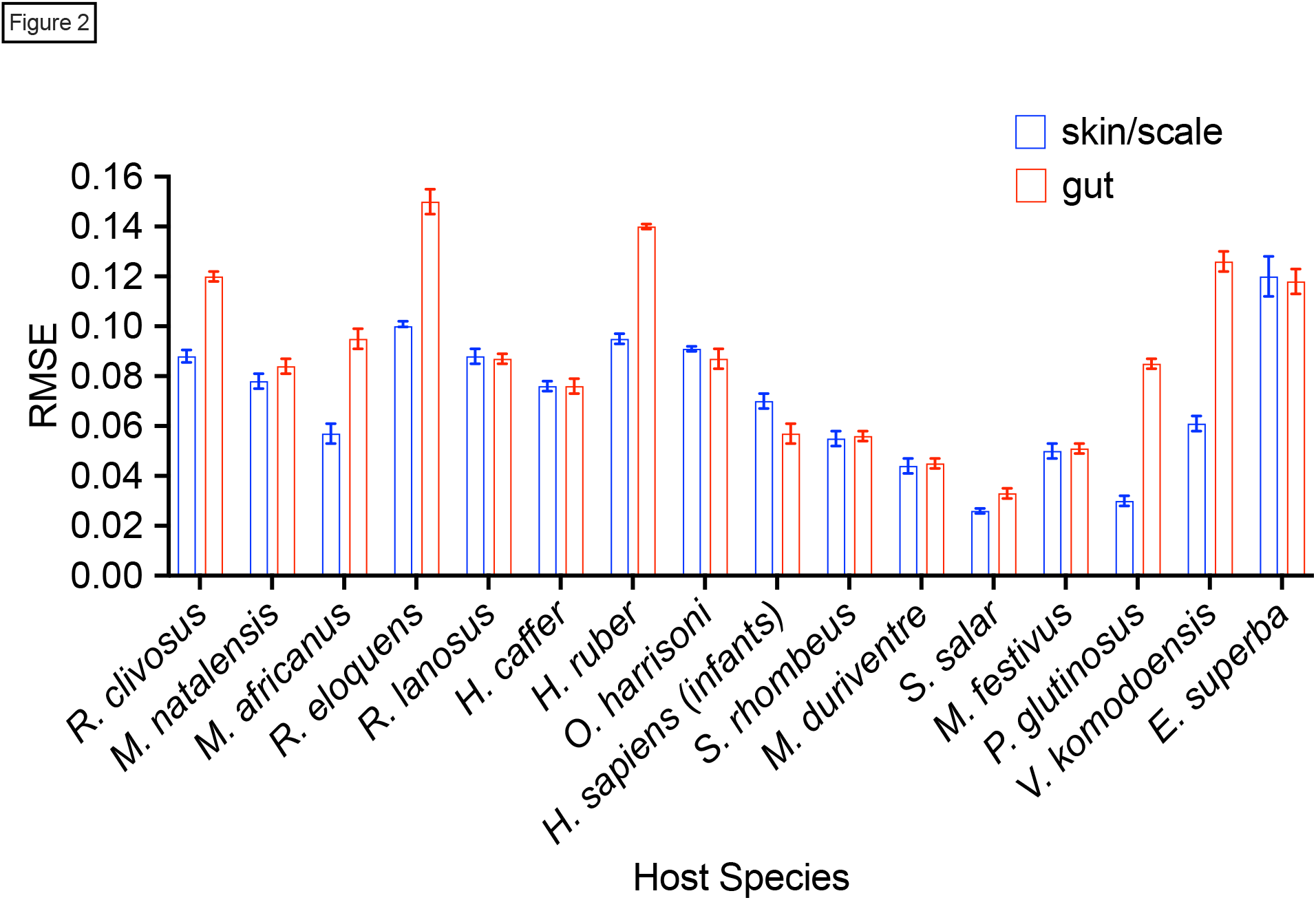
Neutral model fits of empirical microbial communities with the source pool of microbes defined to contain microbes from both tissue types. Error bars represent 95% confidence intervals generated by bootstrapping data.

In this study, we used a modelling-based approach to understand the ecological processes that underlie the assembly of microbiomes associated with gut and skin or scale tissues across a panel of animal hosts. This approach revealed that for the majority of animal species considered, the skin or scale microbiome was more neutrally assembled than the gut microbiome. This may be the result of variation among body sites in susceptibility to microbes dispersing from the environment or host activities that promote/deter microorganisms. We also noted variation among broad host taxonomic groups in the contribution of neutral processes to microbiome assembly among tissue types. Determining the factors that drive these differences (e.g. host lifestyle or immune system complexity) will be an important avenue of future research. Overall, our observations suggest that the gut environment tends to be more selective than the skin or scale environment and showcases the utility of this method to reveal potentially unique features of tissue environments in animal hosts.

## References

1. M. McFall-Ngai, M.G. Hadfield, T.C.G. Bosch, H.V. Carey, T. Domazet-Lošo, A.E. Douglas, N. Dubilier, G. Eberl, T. Fukami, S.F. Gilbert, U. Hentschel, N. King, S. Kjelleberg, A.H. Knoll, N. Kremer, S.K. Mazmanian, J.L. Metcalf, K. Nealson, N.E. Pierce, J.F. Rawls, A. Reid, E.G. Ruby, M. Rumpho, J.G. Sanders, D. Tautz, & J.J. Wernegreen, 2013. Animals in a bacterial world, a new imperative for the life sciences, Proc. Natl. Acad. Sci. U.S.A. 110 (9) 3229–3236, 10.1073/pnas.1218525110 (2013).

2. Broderick, N. A., Buchon, N., & Lemaitre, B. (2014). Microbiota-induced changes in drosophila melanogaster host gene expression and gut morphology. mBio, 5(3), e01117–14. 10.1128/mBio.01117-14

3. Perez Perez, G. I., Gao, Z., Jourdain, R., Ramirez, J., Gany, F., Clavaud, C., Demaude, J., Breton, L., & Blaser, M. J. (2016). Body Site Is a More Determinant Factor than Human Population Diversity in the Healthy Skin Microbiome. PloS one, 11(4), e0151990. 10.1371/journal.pone.0151990

4. Costello, E. K., Carlisle, E. M., Bik, E. M., Morowitz, M. J., & Relman, D. A. (2013). Microbiome assembly across multiple body sites in low-birthweight infants. mBio, 4(6), e00782–13. 10.1128/mBio.00782-13

5. Faust K, Sathirapongsasuti JF, Izard J, Segata N, Gevers D, et al. 2012. Microbial Co-occurrence Relationships in the Human Microbiome. PLOS Computational Biology 8(7): e1002606. 10.1371/journal.pcbi.1002606

6. Asangba AE,,Mugisha L,Rukundo J, Lewis RJ, Halajian A, Cortés-Ortiz L, Junge RE, Irwin MT, Karlson J, Perkin A, Watsa M,Erkenswick G,Bales KL, Patton DL, Jasinska AJ,Fernandez-Duque E, Leigh SR, Stumpf RM. 2022. Large Comparative Analyses of Primate Body Site Microbiomes Indicate that the Oral Microbiome Is Unique among All Body Sites and Conserved among Nonhuman Primates. Microbiol Spectr 10:e01643–21.

7. Vellend M. 2010. Conceptual synthesis in community ecology. The Quarterly review of biology, 85(2), 183–206. 10.1086/652373

8. Burns, A. R., Stephens, W. Z., Stagaman, K., Wong, S., Rawls, J. F., Guillemin, K., & Bohannan, B. J. 2016. Contribution of neutral processes to the assembly of gut microbial communities in the zebrafish over host development. The ISME journal, 10(3), 655–664. 10.1038/ismej.2015.142

9. Bolyen, E., Rideout, J. R., Dillon, M. R., Bokulich, N. A., Abnet, C. C., Al-Ghalith, G. A., Alexander, H., Alm, E. J., Arumugam, M., Asnicar, F., Bai, Y., Bisanz, J. E., Bittinger, K., Brejnrod, A., Brislawn, C. J., Brown, C. T., Callahan, B. J., Caraballo-Rodríguez, A. M., Chase, J., Cope, E. K., … Caporaso, J. G. (2019). Reproducible, interactive, scalable and extensible microbiome data science using QIIME 2. Nature biotechnology, 37(8), 852–857. 10.1038/s41587-019-0209-9

10. Callahan, B. J., McMurdie, P. J., Rosen, M. J., Han, A. W., Johnson, A. J., & Holmes, S. P. 2016. DADA2: High-resolution sample inference from Illumina amplicon data. Nature methods, 13(7), 581–583. 10.1038/nmeth.3869

11. Bokulich, N. A., Kaehler, B. D., Rideout, J. R., Dillon, M., Bolyen, E., Knight, R., Huttley, G. A., & Gregory Caporaso, J. 2018. Optimizing taxonomic classification of marker-gene amplicon sequences with QIIME 2’s q2-feature-classifier plugin. Microbiome, 6(1), 90. 10.1186/s40168-018-0470-z

12. Pedregosa, F., Varoquaux, G., Gramfort, A., Michel, V., Thirion, B., Grisel, O., … & Duchesnay, É. 2011. Scikit-learn: Machine learning in Python. the Journal of machine Learning research, 12, 2825–2830.

13. Quast, C., Pruesse, E., Yilmaz, P., Gerken, J., Schweer, T., Yarza, P., Peplies, J., & Glöckner, F. O. 2013. The SILVA ribosomal RNA gene database project: improved data processing and web-based tools. Nucleic acids research, 41(Database issue), D590–D596. 10.1093/nar/gks1219

14. Oksanen J, Simpson G, Blanchet F, Kindt R, Legendre P, Minchin P, O’Hara R, Solymos P, Stevens M, Szoecs E, Wagner H, Barbour M, Bedward M, Bolker B, Borcard D, Borman T, Carvalho G, Chirico M, De Caceres M, Durand S, Evangelista H, FitzJohn R, Friendly M, Furneaux B, Hannigan G, Hill M, Lahti L, Martino C, McGlinn D, Ouellette M, Ribeiro Cunha E, Smith T, Stier A, Ter Braak C, Weedon J 2022. vegan: Community Ecology Package. R package version 2. 6-4, https://vegandevs.github.io/vegan/.

15. Sloan, W. T., Lunn, M., Woodcock, S., Head, I. M., Nee, S., & Curtis, T. P. 2006. Quantifying the roles of immigration and chance in shaping prokaryote community structure. Environmental microbiology, 8(4), 732–740. 10.1111/j.1462-2920.2005.00956.x

16. Sieber, M., Pita, L., Weiland-Bräuer, N., Dirksen, P., Wang, J., Mortzfeld, B., Franzenburg, S., Schmitz, R. A., Baines, J. F., Fraune, S., Hentschel, U., Schulenburg, H., Bosch, T. C. G., & Traulsen, A. 2019. Neutrality in the Metaorganism. PLoS biology, 17(6), e3000298. 10.1371/journal.pbio.3000298

17. Stephens WZ, Burns AR, Stagaman K, Wong S, Rawls JF, Guillemin K, Bohannan BJ. The composition of the zebrafish intestinal microbial community varies across development. ISME J. 2016 Mar;10(3):644–54. doi: 10.1038/ismej.2015.140. Epub 2015 Sep 4. PMID: 26339860; PMCID: PMC4817687.

18. Jani, A. J., Bushell, J., Arisdakessian, C. G., Belcaid, M., Boiano, D. M., Brown, C., & Knapp, R. A. 2021. The amphibian microbiome exhibits poor resilience following pathogen-induced disturbance. The ISME journal, 15(6), 1628–1640. 10.1038/s41396-020-00875-w

19. Custer, G. F., Bresciani, L., & Dini-Andreote, F. 2022. Ecological and Evolutionary Implications of Microbial Dispersal. Frontiers in microbiology, 13, 855859. 10.3389/fmicb.2022.855859

20. She, J. J., Liu, W. X., Ding, X. M., Guo, G., Han, J., Shi, F. Y., Lau, H. C., Ding, C. G., Xue, W. J., Shi, W., Liu, G. X., Zhang, Z., Hu, C. H., Chen, Y., Wong, C. C., & Yu, J. 2024. Defining the biogeographical map and potential bacterial translocation of microbiome in human ‘surface organs’. Nature communications, 15(1), 427. 10.1038/s41467-024-44720-6

21. Venkataraman, A., Bassis, C. M., Beck, J. M., Young, V. B., Curtis, J. L., Huffnagle, G. B., & Schmidt, T. M. 2015. Application of a neutral community model to assess structuring of the human lung microbiome. mBio, 6(1), e02284–14. 10.1128/mBio.02284-14

