## Supplemental table 1 for "Assembly of skin microbiomes is more neutral than gut microbiomes in multiple animal species"

Supplementary Table 1: Datasets used in this study

| Host | Number of Individuals | Body Sites Sampled | Reference |
| --- | --- | --- | --- |
| *M*y*lossoma duriventre* | 45 | Surface swab midgut, hindgut | Sylvain et al. 2020 PMID **32503908** |
| *Serrasalmus rhombeus* | 33 | Surface swab, midgut, hindgut | Sylvain et al. 2020 PMID **32503908** |
| *Mesonata festivus* | 33 | Surface swab, midgut, hindgut | Sylvain et al. 2020 PMID **32503908** |
| *Varanus komodoensis* | 34 | Feces, Skin | Hyde et al. 2016  PMID **27822543** |
| *Salmo salar* | 74 | Fish surface, whole intestine | Webster et al. 2018  PMID 29915104 |
| *Euphasia superba* | 22 | Moult, feces | Clark et al. 2019  PMID 30697197 |
| *Plethodon glutinosus* | 60 | Surface swab, feces | Walker et al. 2019  PMID **31802185** |
| *Rhinolopus eloquens* | 23 | Multiple body site skin swab (pooled), distal colon | Lutz et al. 2019, PMID **31719140** |
| *Miniopterus natalensis* | 26 | Multiple body site skin swab (pooled), distal colon | Lutz et al. 2019, PMID **31719140** |
| *Rhinolophus clivosus* | 21 | Multiple body site skin swab (pooled), distal colon | Lutz et al. 2019, PMID **31719140** |
| *Hipposideros caffer* | 34 | Multiple body site skin swab (pooled), distal colon | Lutz et al. 2019, PMID **31719140** |
| *Hipposideros ruber* | 20 | Multiple body site skin swab (pooled), distal colon | Lutz et al. 2019, PMID **31719140** |
| *Otomops harrissoni* | 25 | Multiple body site skin swab (pooled), distal colon | Lutz et al. 2019, PMID **31719140** |
| *Miniopterus africanus* | 22 | Multiple body site skin swab (pooled), distal colon | Lutz et al. 2019, PMID **31719140** |
| *Rousettus lanosus* | 31 | Multiple body site skin swab (pooled), distal colon | Lutz et al. 2019, PMID **31719140** |
| *Homo sapiens* | 38 | Multiple body sites (variable), feces | Younge et al. 2018,  PMID 29855335 |
